# Anaerobic Bacteria in the Gut Microbiota Confer Colonization Resistance Against ESBL-Producing *Escherichia coli* in Mice

**DOI:** 10.1101/2025.04.06.647462

**Authors:** Mika Murata, Yoshitomo Morinaga, Daisuke Sasaki, Katsunori Yanagihara

**Author notes:** **Corresponding author** Yoshitomo Morinaga, MD, PhD, Department of Microbiology, Graduate School of Medicine and Pharmaceutical Sciences, University of Toyama, 2630 Sugitani, Toyama 930-0194, Japan. **Authorship statement:** Conceptualization: YM; Methodology: MM, YM, and DS; Validation: MM and DS; Formal Analysis: MM and DS; Investigation: MM, YM, and DS; Resources: N/A; Data Curation: YM and DS; Writing–Original Draft Preparation: MM and YM; Writing–Review and Editing: MM, YM, DS, and KY; Visualization: MM and YM; Supervision: YM and KY; Project Administration: YM; Funding acquisition: MM, YM, and KY. All authors meet the ICMJE authorship criteria.

## Abstract

Antimicrobial resistance (AMR) is a growing health concern worldwide, and gut microbiota play a significant role in its spread. This study aimed to investigate the impact of antibiotic-induced alterations in gut microbiota on the colonization of extended-spectrum β-lactamase (ESBL)-producing *Escherichia coli* in a mouse model. C57BL/6J mice were treated with various antibiotics (ampicillin, vancomycin, neomycin, metronidazole, or a cocktail of all four) prior to oral inoculation with ESBL-producing *E. coli*. 16S rRNA metagenomics analysis revealed significant alterations in gut microbiota composition and diversity following antibiotic treatment. Notably, ampicillin, vancomycin, and the antibiotic cocktail dramatically increased colonization by ESBL-producing *E. coli*, whereas metronidazole and neomycin treatments had minimal effects. Linear discriminant analysis highlighted that specific anaerobic bacterial groups, namely *Bacteroidales*, Lachnospiraceae, and Ruminococcaceae, were inversely correlated with colonization by ESBL-producing *E. coli*. Collectively, these findings suggest that diverse anaerobic bacteria play a crucial role in resistance against AMR bacteria colonization. This study provides insights into the complex interactions between gut microbiota and AMR colonization that could aid in the development of future strategies for risk assessment and eradication of drug-resistant bacteria.

## INTRODUCTION

The recent increase in antimicrobial resistant (AMR) bacteria is a significant global concern. The estimated number of deaths annually caused by AMR infections could reach approximately 10 million by 2050 if no effective approaches are implemented [1]. Notably, AMR among gut-resident bacteria such as *Escherichia coli, Klebsiella* spp., and *Enterobacter* spp. has been rapidly increasing globally [2-5]. However, there is limited evidence describing the mechanisms through which AMR among these bacteria has increased and spread among humans.

The intestinal environment serves as an AMR reservoir that is favorable for the transfer of AMR-related genes through mechanisms such as horizontal gene transfer [6-9]. The introduction of AMR bacteria into the intestinal environment is the first step for the colonization. Clinical research has revealed that patients receiving antibiotic therapy have an elevated risk of AMR gut colonization, often associated with antibiotic-induced dysbiosis, which disrupts the composition and function of the gut microbiota [10, 11]. *In vivo* experiments have revealed that dysbiosis elevates the amount of AMR bacteria in feces [12]. However, while some insights have been gained into the role of the intestinal environment in AMR, many details remain unclear and are still being investigated.

A healthy gut microbiota consists of a complex and diverse community that is resistant to pathogen invasion [13]. However, antibiotics disrupt bacterial community balance, enabling the invasion of pathogens such as *Clostridioides difficile* and *Salmonella* spp. [14, 15]. This protective function is commonly referred to as colonization resistance, which involves direct and indirect competitive interactions between pathogens and commensal bacteria [13, 16]. However, unlike these pathogens, AMR bacteria from the family Enterobacteriaceae are known to be carriers and coexist with the commensal bacteria in the body [4].

In this study, to reveal the specific bacteria that contribute to or protect from colonization, microbial signatures associated with resistance to AMR bacteria colonization were analyzed in a disrupted gut microbiota in mice.

## MATERIALS AND METHODS

### Animals

Four- to six-week-old female C57BL/6J mice were purchased from Charles River Laboratories Japan, Inc. (Kanagawa, Japan). All animals were housed in a pathogen-free environment at the Laboratory Animal Center for Biomedical Science at Nagasaki University and were provided with sterile food and water. The mice were co-housed for at least two weeks prior to the experiments. The Ethics Review Committee for Animal Experimentation (Institutional Animal Care and Use Committee (IACUC) of Nagasaki University) approved all the experimental protocols used in this study (Protocol Number: 1503101199).

### Mouse treatment and gut colonization model

After co-housing, the C57BL/6J mice were administered regular or antibiotic-containing water (1 g/L ampicillin [TCI Chemicals, Tokyo, Japan], 0.5 g/L vancomycin [Fujifilm Wako Chemicals, Osaka, Japan], 1 g/L neomycin [TCI Chemicals], or 1 g/L metronidazole [TCI Chemicals], or a cocktail containing ampicillin, vancomycin, neomycin, and metronidazole at the same concentrations) via their drinking water for 3 days. Fecal pellets were collected for 16S rRNA metagenome analysis.

To induce the colonization of extended-spectrum β-lactamase (ESBL)-producing *E. coli* in the gut, following an additional 2-day interval where the mice received regular water to wash out the residual antibiotics, the mice were orally inoculated with *E. coli* N001, a clinical isolate of sequence type 131 carrying CTX-M-15-type ESBL, at Nagasaki University Hospital, at 2 × 10^7^ CFU/mouse.

### Quantification of bacteria in feces

Fecal pellets were collected from each mouse and dissolved in DPBS. The fecal samples were plated on MacConkey agar (Becton, Dickinson and Company, Sparks, MD) with 50 mg/L cefotaxime. Because ESBL-producing *E. coli* can grow on the plate, the number of colonies was determined to be the number of AMR bacteria.

### PCR amplification and 16S rRNA gene sequencing preparation

DNA was extracted using a Quick-DNA Fecal/Soil Microbe Miniprep Kit (ZYMO Research, Irvine, CA), according to the manufacturer’s instructions. The V1-V2 region of the bacterial 16S rRNA genes was amplified, and sequencing was performed using the Ion Torrent Personal Genome Analyzer, as previously described[17]. Briefly, the PCR amplicons were purified using an AMPure XP Kit (Beckman Coulter, Indianapolis, IN), and the purified amplicons were mixed in equimolar amounts. The samples were adjusted to a final concentration of 100 pM for use as template DNA in emulsion PCR. Emulsion PCR and enrichment were performed using an Ion PGM HiQ View OT2 Kit (Thermo Fisher Scientific, Waltham, MA) according to the manufacturer’s instructions. The enriched samples were loaded onto an Ion 318 chip, and sequencing was performed using the Ion Torrent Personal Genome Analyzer with an Ion PGM HiQ View Sequencing Kit (Thermo Fisher Scientific, Waltham, MA), according to the manufacturer’s instructions.

### Sequence analysis

The sequencing reads were analyzed using CLC Genomics Workbench version 12.0.1 and CLC Microbial Genomics Module version 3.6.11 (QIAGEN N.V., Venlo, Netherlands), as previously described [17]. Briefly, the primer sequences were removed, and the read length was trimmed between 100 bp and 400 bp under a 0.05% quality limit. Samples with fewer than 100 reads and those with less than 50% of the median read count were excluded from the further analyses. Chimeric reads were detected and filtered using the chimera crossover detection algorithm with default parameters. The reads were categorized into operational taxonomic units (OTUs) with 97% similarity, and taxonomy was assigned using SILVA release 132. The OTUs were aligned using MUSCLE software integrated into CLC software. The number of OTUs, Shannon index (alpha diversity), and Weighted UniFrac distances were determined using the CLC Microbial Genomics Module.

### Linear discriminant analysis (LDA)

Differential microbial abundance was analyzed using Linear Discriminant Analysis Effect Size (LefSe) on the Galaxy online application (http://galaxy.biobakery.org/). The LDA log score threshold was set to 2.0, followed by the Kruskal–Wallis test and a Wilcoxon test with a cut-off of p□<□0.05.

### Statistical analysis

Groups were compared using the Mann-Whitney test with Prism version 8 (GraphPad software, CA). Statistical significance (alpha level) was set as p ≤ 0.05. To compare the beta diversity, the data were analyzed using PERMANOVA analysis in CLC, and statistical significance (alpha level) was set at p ≤ 0.05, using the false discovery rate method.

## RESULTS

### Antibiotic exposure induces alterations in gut microbiota

To determine the characteristics of the altered gut microbiota, mice were orally challenged with four types of antibiotics or a cocktail of all four antibiotics via their drinking water, and 16S rRNA metagenome was analyzed in fecal samples (Fig. 1A). The Shannon index, which indicates alpha-diversity, of the fecal microbiota was significantly decreased in the ampicillin-, vancomycin-, metronidazole-, and antibiotic cocktail-treated mice, but not in neomycin-treated mice, compared to the control mice (Fig. 1B). However, all antibiotic-treated mice exhibited significant differences in the beta-diversity compared to the control mice (Fig. 1C, p < 0.05 compared with the control group).

**Figure 1.**
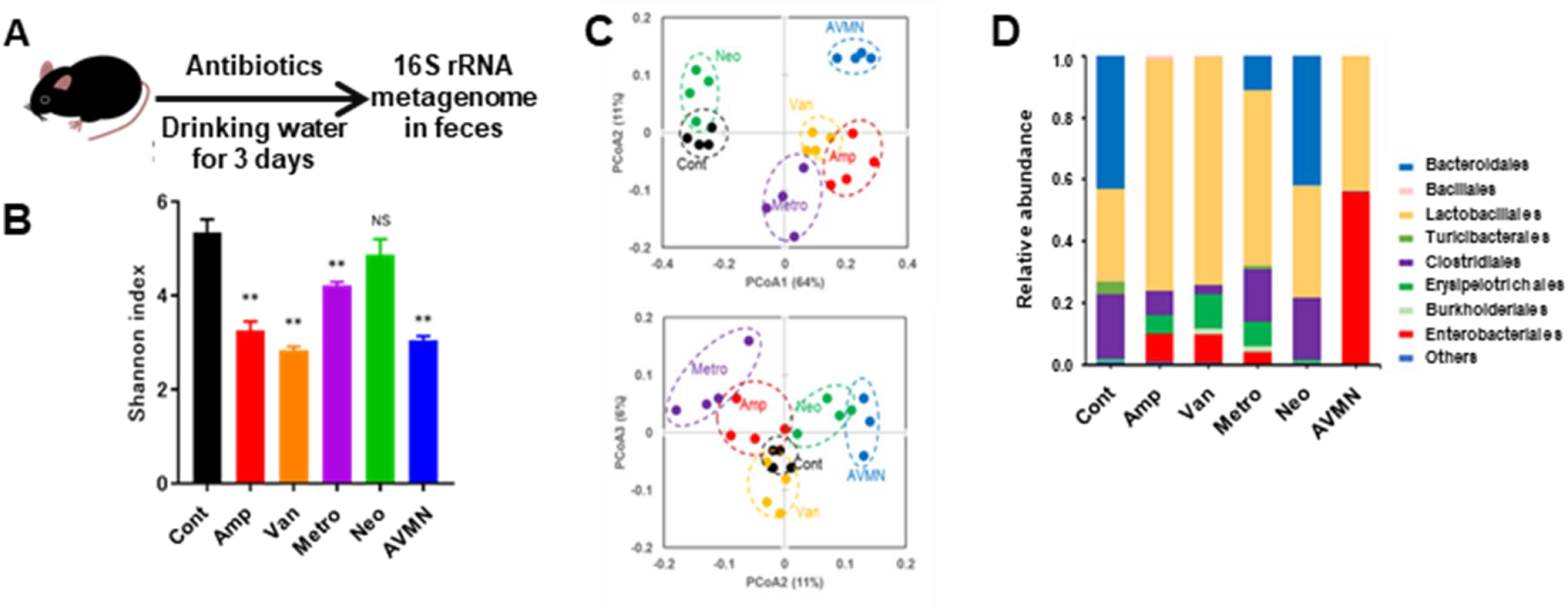
Characteristics of the gut microbiota following antibiotic treatment. **(A)** A schematic illustration of the experimental timeline. **(B)** The alpha diversity (Shannon index) of the gut microbiota following antibiotic treatment (n = 4, each). **(C)** Beta diversity (Weighted Unifrac) of the gut microbiota following antibiotic treatment. Each dot represents an individual mouse. **(D)** The composition of the gut microbiota after treatment with antibiotics. Data are expressed as the mean of four mice from each group. Cont: control, Amp: ampicillin, Van: vancomycin, Metro: metronidazole, Neo: neomycin, AVMN: a cocktail of all four antibiotics. Error bars represent the standard error of the mean. ** indicates p < 0.01 compared to the control.

The gut microbiota were analyzed, revealing that Bacteroidetes, Lactobacillales, and Clostridiales were predominant in the control mice (Fig. 1D). In contrast, in ampicillin-, vancomycin-, or neomycin-treated mice, the proportions of Lactobacillales, Erysipelotrichales, and Enterobacteriales were elevated, while those of Clostridiales and Bacteroidales were decreased. The gut microbiota composition of the neomycin-treated mice was most similar to that of the control mice; however, Turicibacteriales were rarely observed. In antibiotic cocktail-treated mice, the major bacteria were Enterobacteriales and Lactobacillales.

### Identifying bacterial signatures associated with colonization resistance and permissiveness in antibiotic-treated mice

Following this, to confirm the phenotype of AMR colonization during the early phase following inoculation in antibiotic-induced altered gut microbiota, mice were orally inoculated with ESBL-producing *E. coli* (Fig. 2A). The number of bacteria in feces was quantified (Fig. 2B), and the number of AMR bacteria in feces after 24 h was 4.3 ± 0.2 Log_10_CFU/g feces (mean ± SEM) in the control mice. Meanwhile, in metronidazole- and neomycin-treated mice, the counts were 4.9 ± 0.2 Log_10_CFU/g feces and 4.6 ± 0.2 Log_10_CFU/g feces, respectively. In contrast, the number of AMR bacteria was markedly elevated in the ampicillin-, vancomycin-, and antibiotic cocktail-treated mice at 7.0 ± 0.1 Log_10_CFU/g feces, 6.9 ± 0.1 Log_10_CFU/g feces, and 7.3 ± 0.1 Log_10_CFU/g feces, respectively.

**Figure 2.**
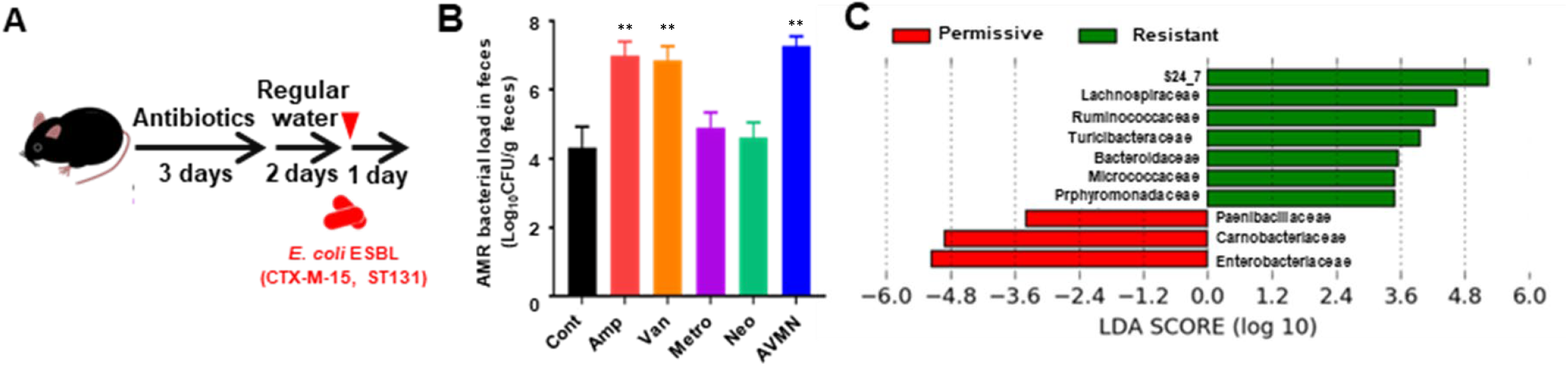
Bacterial signatures of colonization resistance during the early phase of ESBL-producing *E. coli* inoculation. **(A)** A schematic illustration of the experimental time course. **(B)** AMR bacterial loads in feces 24 h after inoculation of ESBL-producing *E. coli* (n = 4). **(C)** Linear discriminant analysis between the colonization-permissive group (ampicillin-, vancomycin-, and antibiotic cocktail-treated mice) and colonization-resistant group (control, neomycin-, and metronidazole-treated mice). Cont: control, Amp: ampicillin, Van: vancomycin, Metro: metronidazole, Neo: neomycin, AVMN: a mix of all four antibiotics. Error bars represent the standard error of the mean. ** indicates p < 0.01 compared to the control.

Finally, to characterize the representative bacteria associated with colonization resistance, LDA was performed (Fig. 2C). The mice groups were categorized into either a colonization-resistant group (control, neomycin-, and metronidazole-treated mice) or a colonization-permissive group (ampicillin-, vancomycin-, and antibiotic cocktail-treated mice). LDA revealed that Bacteroidales, Lachnospiraceae, and Ruminococcaceae were dominant in the colonization-resistant group, whereas Enterobacteriaceae and Carnobacteriaceae were dominant in the colonization-permissive group.

## DISCUSSION

The present study demonstrated that alterations in gut microbiota affect the quantity of ESBL-producing *E. coli* during the early phase of its introduction into the intestinal environment. Moreover, the abundance of anaerobic bacteria was inversely correlated with the quantity of ESBL-producing *E. coli* in the gut.

Similar to previous studies in mice and humans [18, 19], antibiotic treatment induced dysbiosis in the gut microbiota. However, in this present study, not all antibiotics promoted the outgrowth of ESBL-producing *E. coli* in the gut compared to the control group, and no outgrowth was observed following treatment with metronidazole or neomycin. Similar findings were also observed in a previous study on *Klebsiella pneumoniae* [20]. These results indicate that, at least during the initial stages of AMR colonization in the intestinal environment, the disruption of specific bacterial groups may play a more significant role than the dysbiosis of the gut microbiota.

Regarding the specific bacterial groups, the present study identified unique anaerobic bacterial groups, including S24_7, Lachnospiraceae, and Ruminococcaceae, that were inversely correlated with ESBL-producing *E. coli* colonization. S24_7, now known as Muribaculaceae, is a family of bacteria within the order Bacteroidales. Similar to our findings, anaerobic bacteria from these groups have been demonstrated to have an inverse correlation to the carriage of drug-resistant *Klebsiella pneumoniae* in mice [21, 22]. Members of Lachnospiraceae and Ruminococcaceae have also been recognized as characteristic of the human microbiota, which exhibits less colonization with drug-resistant Gram-negative bacteria [23, 24]. Thus, the anaerobic bacteria involved in colonization resistance may be similar among drug-resistant Enterobacteriaceae species.

Our findings could aid in establishing risk assessment methods for detecting drug-resistant bacteria and eradication strategies based on the characteristics of the microbiota. Although there could be differences in the bacterial communities and their compositions between humans and mice, over 80% of annotatable functions have been found to be shared between the microbiota of the two [25]. Therefore, it is crucial to consider the characteristic functions of bacteria. For example, the families Lachnospiraceae and Ruminococcaceae are major producers of butyrate in the gut, which can inhibit the growth of carbapenem-resistant *E. coli* by exploiting its fitness disadvantages [26]. The genus *Bacteroides* has been demonstrated to be associated with microbiota recovery following antibiotic-induced dysbiosis [27]. Furthermore, short-chain fatty acids such as butyrate produced by the gut microbiota have been demonstrated to inhibit Enterobacteriaceae but have no inhibitory effect on Bacteroides spp. [28]. Therefore, for the clinical application involving resistance to the colonization of drug-resistant bacteria, it may be effective to employ experimental animals as models, focusing on commensal bacteria that share characteristic functions with humans, such as metabolic products.

However, this study has several limitations. Firstly, because we only evaluated ESBL-producing *E. coli*, it is vital to investigate whether similar findings are observed for other important AMR bacterial strains such as carbapenemase-producing Enterobacteriaceae and vancomycin-resistant Enterococcus. Second, long-term colonization was not evaluated in the present study, which requires further validation.

In conclusion, anaerobic bacteria, including the order Bacteroidales and the families Lachnospiraceae and Ruminococcaceae, were identified as key bacterial groups that influence initial intestinal colonization following exposure to ESBL-producing *E. coli* and were strongly correlated with colonization resistance.

## TRANSPARENCY DECLARATION

### Data availability

The authors confirm that data supporting the findings of this study are available within the article.

### Conflicts of interest

The authors have no conflicts of interest to declare.

### Funding

This study was supported by the Japanese Society of Laboratory Medicine Fund for the Promotion of Scientific Research (MM), the Research Program on Emerging and Re-emerging Infectious Diseases from the AMED (grant number JP20fk0108133 (YM and KY) and grant number JP22fk0108560 (YM)), and Japan Society for the Promotion of Science (JSPS) KAKENHI grant number JP20K08821 (YM). The funding bodies played no role in the design of the study, collection, analysis, or interpretation of data, nor in writing the manuscript.

## Notes

### Competing Interest Statement

The authors have declared no competing interest.

